# The m^6^A reader protein YTHDF2 facilitates HTLV-1 infectious and mitotic propagation by stabilizing Tax RNA

**DOI:** 10.64898/2025.12.05.692549

**Authors:** Yi Liang, Chenxin Tan, Xinyu Chang, Chenyang Lyu, Dongmei Liu, Shuwen Xu, Dian Zhu, Benquan Liu, Xiaoyi Yuan, Xiaorui Zuo, Xiao-Min Liu, Guangyong Ma

## Abstract

N6-methyladenosine (m^6^A) is one of the important RNA modifications that affect RNA abundance and function and has also been implicated in viral infection. In this study, we identify a number of m^6^A modification sites within human T-cell leukemia virus type 1 (HTLV-1) RNAs, particularly in the RNA of the Tax oncogene. We demonstrate that YTH domain family, member 2 (YTHDF2), a key m^6^A reader protein that binds m^6^A-modified RNAs and influences RNA metabolism, is required for HTLV-1 replication in both *de novo* and persistently infected cells. Mechanistically, YTHDF2 interacts with and stabilizes Tax RNA in an m^6^A-dependent manner, thereby promoting Tax-driven HTLV-1 replication. Importantly, YTHDF2 activates oncogenic cellular pathways and promotes proliferation in HTLV-1 infected cells, aligning with the functions of Tax. Overall, our findings characterize YTHDF2 as a host factor critical to HTLV-1 RNA metabolism and viral propagation, offering novel insights into the understanding of HTLV-1 life cycle and the development of targeted interventions.

**Importance:** m^6^A is an RNA modification that plays crucial roles in physiological and pathological conditions; however, its roles in HTLV-1 infection are poorly understood. In this study, we identify that m^6^A modification is widely present in HTLV-1 RNAs including Tax. We demonstrate that YTHDF2, a prominent m^6^A reader protein, enhances the stability of Tax RNA, thereby promoting both *de novo* and persistent HTLV-1 replication. Our findings position YTHDF2 as an essential factor for HTLV-1 persistence and suggest it as a potential therapeutic target for viral clearance.

## Introduction

HTLV-1 is an oncogenic human retrovirus that infects 5-10 million people worldwide(1) and causes an aggressive blood cancer known as adult T-cell leukemia/lymphoma (ATLL)(2). HTLV-1 replication relies on its transactivator protein Tax to activate viral gene transcription, thereby initiating an infectious stage of viral propagation(3). On the other hand, Tax can also promote the proliferation and transformation of HTLV-1 infected CD4 T cells, leading to the mitotic propagation of the virus in a passive manner(4). Accordingly, Tax has been considered a key viral factor responsible for HTLV-1 replication and pathogenesis. Although the role of Tax is challenged by evidence that Tax expression is often silenced in ATLL patients(5–11), recent observations revealing an inducible or intermittent expression pattern of Tax highlight its essential role in HTLV-1 infection(12–15).

RNA modification regulates the metabolism of RNA in a post-transcriptional manner(16, 17). Among numerous chemical modifications identified on cellular RNA, m^6^A is the most prevalent one in mammals(18). m^6^A exerts its function mainly through the binding of a family of reader proteins to methylated transcripts, which include YTH domain family, member 1 (YTHDF1), YTHDF2, and YTHDF3, and YTH domain-containing protein 1 (YTHDC1) and YTHDC2(19). The binding of these reader proteins to m^6^A-modified mRNAs predominantly regulate RNA stability and translation, resulting in various physiologically relevant processes(20, 21). Very recently, m^6^A reader proteins have been shown to regulate HTLV-1 RNA transcription(22). King et al reported that YTHDF1 suppresses HTLV-1 Tax but increases HBZ expression, and YTHDC1 promotes the nuclear export of Tax RNA(22). Here we performed methylated RNA immunoprecipitation (MeRIP) -qPCR and identified widely present m^6^A modifications in HTLV-1 RNAs, agreeing with the findings by King et al. On the other hand, we focused on YTHDF2 and demonstrated that it is able to bind and stabilize Tax RNA, thereby promoting HTLV-1 infectivity. Our results for the first time systemically characterized the roles of YTHDF2 in HTLV-1, underscoring an essential requirement of YTHDF2 for HTLV-1 infection.

## Results

### Endogenous YTHDF2 is required for HTLV-1 replication

We utilized a frequently used m^6^A modification prediction tool named SRAMP(23) and discovered 19 high-confidence m^6^A sites across the complete HTLV-1 genomic RNA (Fig. S1). By performing MeRIP-qPCR, we confirmed the presence of these m^6^A sites in HeLa S3 cells *de novo* infected with an HTLV-1 infectious clone, pX1MT-M(24) (Fig. 1A and B). Furthermore, m^6^A modifications were also observed in ATL-T, a T cell line persistently infected with HTLV-1 (Fig. 1B). It is worth noting that the m^6^A sites appear more abundant in the pX region of HTLV-1 (Fig. 1B), in particular sequences encoding Tax (p7578 fragment in Fig. 1B). We continued to identify by MeRIP-qPCR that Tax RNA was indeed modified with m^6^A in both HTLV-1 *de novo* and persistently infected cells (Fig. 1C).

**Fig. 1.**
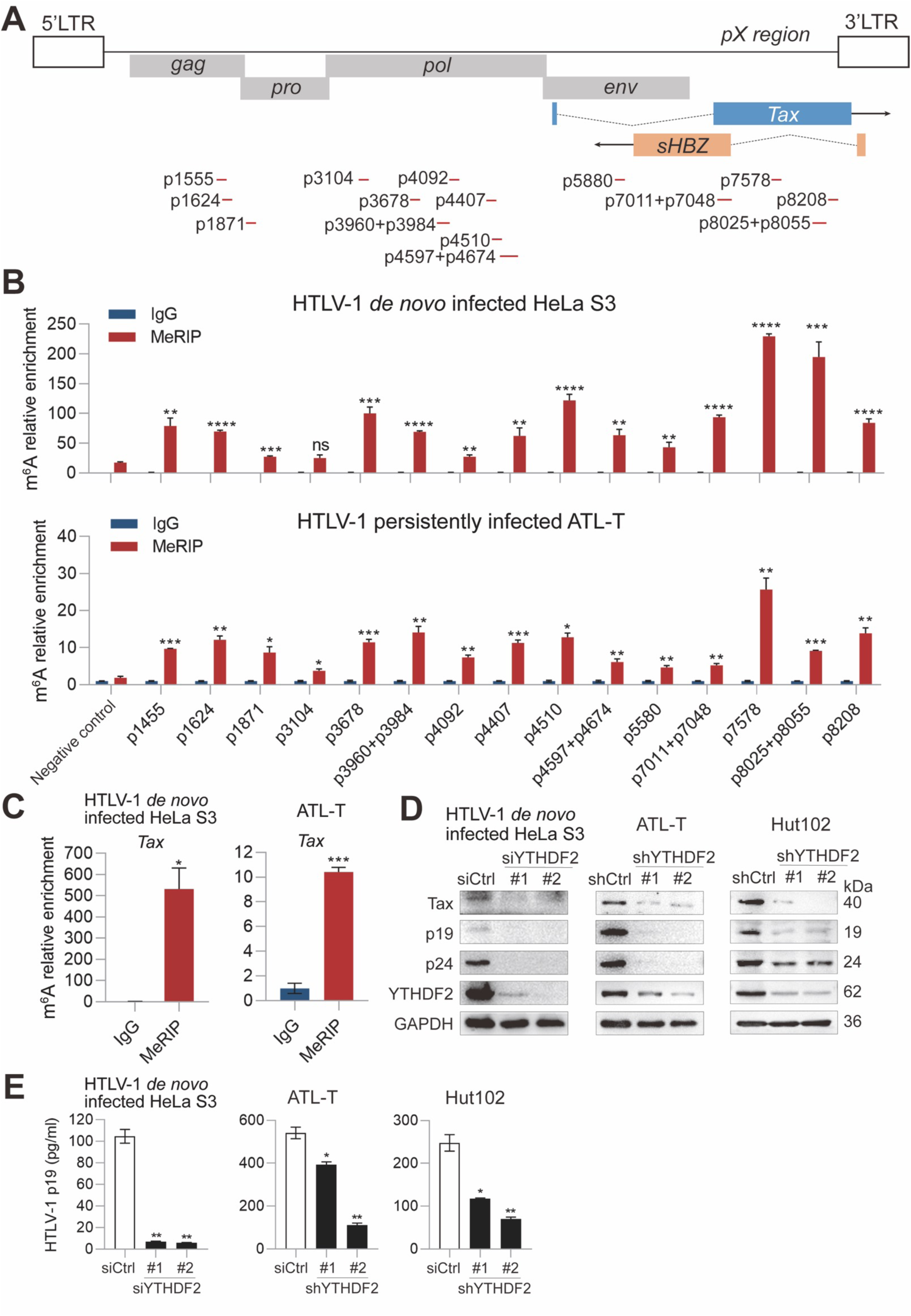
Endogenous YTHDF2 is required for HTLV-1 replication. (A) (*Upper*) Schematic diagram of the HTLV-1 genome structure; (*Lower*) Design of 15 MeRIP-qPCR primer pairs (Table S1) covering 19 high-confidence sites. The red lines indicate the amplification regions. (B) MeRIP-qPCR analysis detected m^6^A modifications of HTLV-1 RNAs in HTLV-1 *de novo* infected HeLa S3 (*Upper*) and persistently infected ATL-T cells (*Lower*). (C) MeRIP-qPCR analysis detected m^6^A modifications of Tax RNA in HTLV-1 infected HeLa S3 and ATL-T cells. (D) Immunoblot analysis of HTLV-1 genes and YTHDF2 expression levels in HTLV-1 infected HeLa S3, ATL-T, and Hut102 cells following YTHDF2 knockdown, with GAPDH used for normalization. (E) ELISA analysis of HTLV-1 p19 levels in the supernatants from HTLV-1 infected HeLa S3, ATL-T and Hut102 cells following YTHDF2 knockdown. *P* value was calculated using a two-tailed unpaired Student’s t-test. ns, not significant; *p<0.05, **p<0.01, ***p<0.001, ****p<0.0001. The results are representatives of three independent experiments.

m^6^A modifications require reader proteins to translate the marks into various functions(21, 22). YTHDF2 is an extensively studied m^6^A reader protein and is also implicated in the regulation of viral RNA expression(25); however, its function in HTLV-1 infection remains undefined. By knocking down endogenous YTHDF2, we observed significantly decreased expressions of Tax RNA (Fig. S2A-C) and protein (Fig. 1D) in HTLV-1 *de novo* infected HeLa S3 and persistently infected cell lines ATL-T and Hut102. In addition, the expression of HTLV-1 structural protein Gag was also reduced upon YTHDF2 knockdown at both RNA (Fig. S2A-C) and protein levels (Gag polyprotein is cleaved into the p19 matrix and p24 capsid proteins) (Fig. 1D). Consistently, HTLV-1 virion production measured by the p19 ELISA was suppressed by YTHDF2 knockdown as well (Fig. 1E). These findings suggest that YTHDF2 is required for HTLV-1 replication in both *de novo* and persistently infected cells.

### YTHDF2 stabilizes Tax RNA in an m^6^A-dependent manner

YTHDF2 is able to bind methylated RNAs and regulate their stability(26). By conducting RNA immunoprecipitation (RIP) assays using an anti-YTHDF2 antibody, we successfully demonstrated the interactions between YTHDF2 protein and Tax or Gag RNA at endogenous levels in HTLV-1 infected cell lines (Fig. 2A). Furthermore, the half-lives of Tax and Gag RNAs were significantly reduced when endogenous YTHDF2 was depleted (Fig. 2B and C). To examine whether YTHDF2 exerts an m^6^A-dependent regulation of Tax RNA expression, we generated a Tax-m^6^A-Mut construct which has a single A to C mutation in one of the predicted m^6^A sites (Fig. 2D). Compared to wild-type Tax (Tax-WT), expression of the Tax-m^6^A-Mut construct in HeLa S3 yielded reduced Tax RNA and protein abundances (Fig. 2E and F), confirming the critical role of m^6^A modification in maintaining Tax expression. Whereas knocking down YTHDF2 greatly impaired Tax RNA and protein abundances in Tax-WT transfected HeLa S3, such effect disappeared in the case of Tax-m^6^A-Mut (Fig. 2E and F). Overall, these findings underscore an m^6^A-dependent regulation of Tax RNA by YTHDF2 and stress its positive role in HTLV-1 infection.

**Fig. 2.**
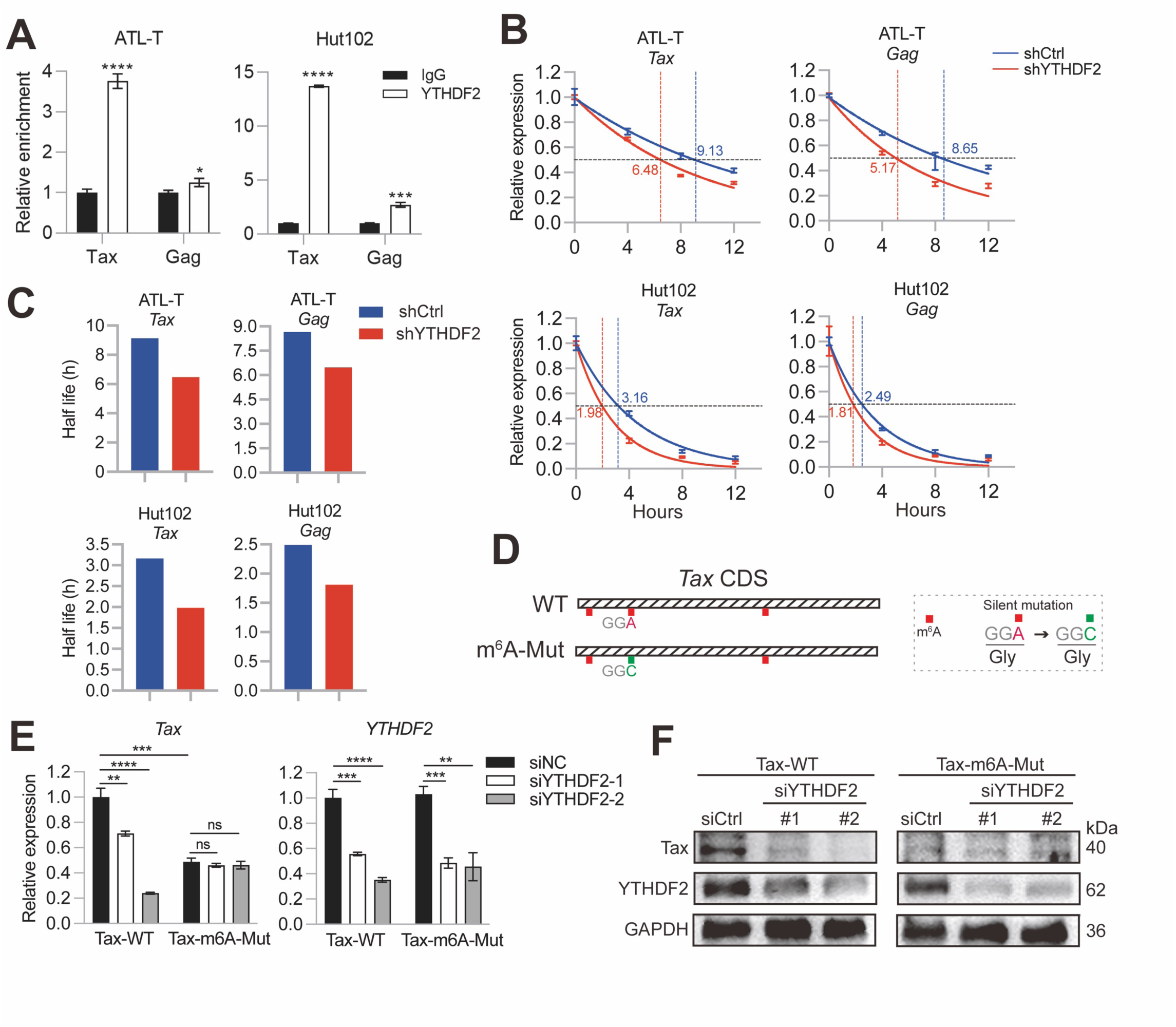
YTHDF2 stabilizes Tax RNA in an m^6^A-dependent manner. (A) RIP-qPCR detection of the interaction between YTHDF2 protein and Tax or Gag RNA in ATL-T and Hut102 cells. (B) The impact of YTHDF2 knockdown (shYTHDF2-1) on the half-lives of Gag and Tax RNAs in ATL-T and Hut102 cells. (C) A bar chart illustrating the half-life of each group depicted in (B). (D) Schematic of constructing Tax m^6^A mutants. A silent mutation (A to C) was introduced at the second m^6^A site. (E) qPCR analysis of the effect of YTHDF2 knockdown on Tax RNA expression in Tax-WT and Tax-m^6^A-Mut. (F) Immunoblot analysis of the expression of Tax protein in Tax-WT and Tax-m^6^A-Mut, upon YTHDF2 knockdown. *P* value was calculated using a two-tailed unpaired Student’s t-test. ns, not significant;*p<0.05, **p<0.01, ***p<0.001, ****p<0.0001. The results are representatives of three independent experiments.

### Endogenous YTHDF2 is required for the proliferation of HTLV-1 infected cells

Tax has been shown to be required for the mitotic growth of HTLV-1 infected cells(3, 15, 27). Since YTHDF2 promotes Tax expression, we hypothesized it might also support the growth of HTLV-1 infected cells. Indeed, knockdown of YTHDF2 in ATL-T and Hut102 (Fig. 3A) resulted in markedly impaired cellular proliferation (Fig. 3B), accompanied by the significantly increased apoptosis (Fig. 3C and D). In contrast, YTHDF2 knockdown did not apparently affect the growth or apoptosis of HTLV-1 uninfected T-cell lines including Jurkat and Hut78 (Fig. S3A-F). Moreover, we established a xenograft mouse tumor model by subcutaneously injecting Hut102, and demonstrated that YTHDF2 knockdown significantly suppressed the proliferation and tumorigenesis of HTLV-1 infected cells *in vivo* (Fig. 3E-G). Therefore, endogenous YTHDF2 appears to be required for Tax-dependent HTLV-1 mitotic propagation both *in vitro* and *in vivo*.

**Fig. 3.**
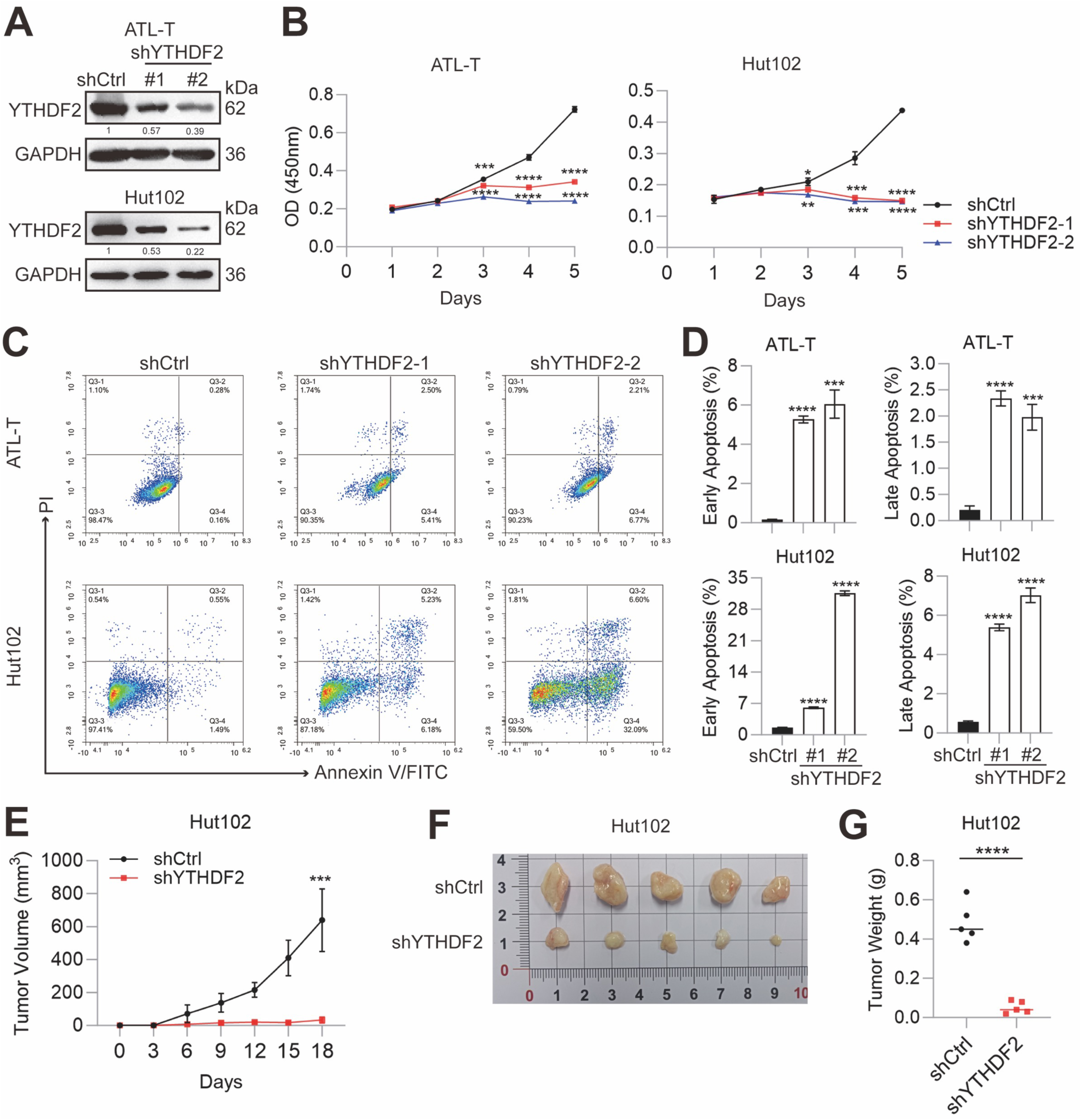
Endogenous YTHDF2 is required for the proliferation of HTLV-1 infected cells. (A) Immunoblot analysis of YTHDF2 knockdown efficiency in (*Upper*) ATL-T and (*Lower*) Hut102 cells, with GAPDH used for normalization. (B) CCK-8 assay examination of cell proliferation in ATL-T and Hut102 after YTHDF2 knockdown. (C) Flow cytometry analysis of apoptosis in (*Upper*) ATL-T and (*Lower*) Hut102 cells after YTHDF2 knockdown. (D) Statistical analysis of early and late apoptosis based on the data in (C). (E-G) The impact of YTHDF2 knockdown (shYTHDF2-1) on subcutaneous tumor growth in xenograft mice established with Hut102 (n=5): (E) Tumor growth curve, (F) Tumors images, and (G) Statistical analysis of tumor weights. *P* value was calculated using a two-tailed unpaired Student’s t-test. *p<0.05, **p<0.01, ***p<0.001, ****p<0.0001. The results are representatives of three independent experiments.

### The YTHDF2–m^6^A–Tax axis is critical to HTLV-1 mitotic propagation

To examine more closely the effect of YTHDF2 on HTLV-1 mitotic propagation, we analyzed the transcriptome changes upon YTHDF2 knockdown in ATL-T and Hut102 (Fig. S4A). Consistent with its impact on Tax expression (Fig. 1), YTHDF2 knockdown resulted in a decrease of global HTLV-1 transcription (Fig. 4A). In addition, endogenous YTHDF2 also profoundly regulates host cell transcription in ATL-T and Hut102, affecting thousands of genes (Fig. 4B). Intriguingly, Gene Set Enrichment Analysis (GSEA) identified that YTHDF2 was significantly associated with multiple oncogenic signature gene sets including MYC Target V1, G2M Checkpoint, MYC Target V2, and Oxidative Phosphorylation in both ATL-T and Hut102 (Fig. 4C and Fig. S4B). We further revealed that YTHDF2 and Tax overlapped in regulating the above oncogenic cellular pathways (Fig. 4D and Fig. S4C), based on the acquisition and subsequent analysis of the Tax-targeted transcriptome in HeLa S3 cells ectopically expressing the wild type or Tax-knockout (KO) pX1MT-M(24). In support of this notion, Kyoto Encyclopedia of Genes and Genomes (KEGG) analysis on YTHDF2-targeted transcriptome in ATL-T and Hut102 both enriched the HTLV-1 infection pathway (Fig. 4E). To provide further evidence for the orchestration of HTLV-1 mitotic propagation by the YTHDF2–m^6^A–Tax axis, a rescue experiment was performed: we enforced Tax expression in YTHDF2 knockdown ATL-T and Hut102 (Fig. 5A and B), and successfully rescued the impaired cell viability caused by YTHDF2 depletion (Fig. 5C). These results likely pinpoint YTHDF2 as a crucial factor regulating HTLV-1 persistent infection via Tax.

**Fig. 4.**
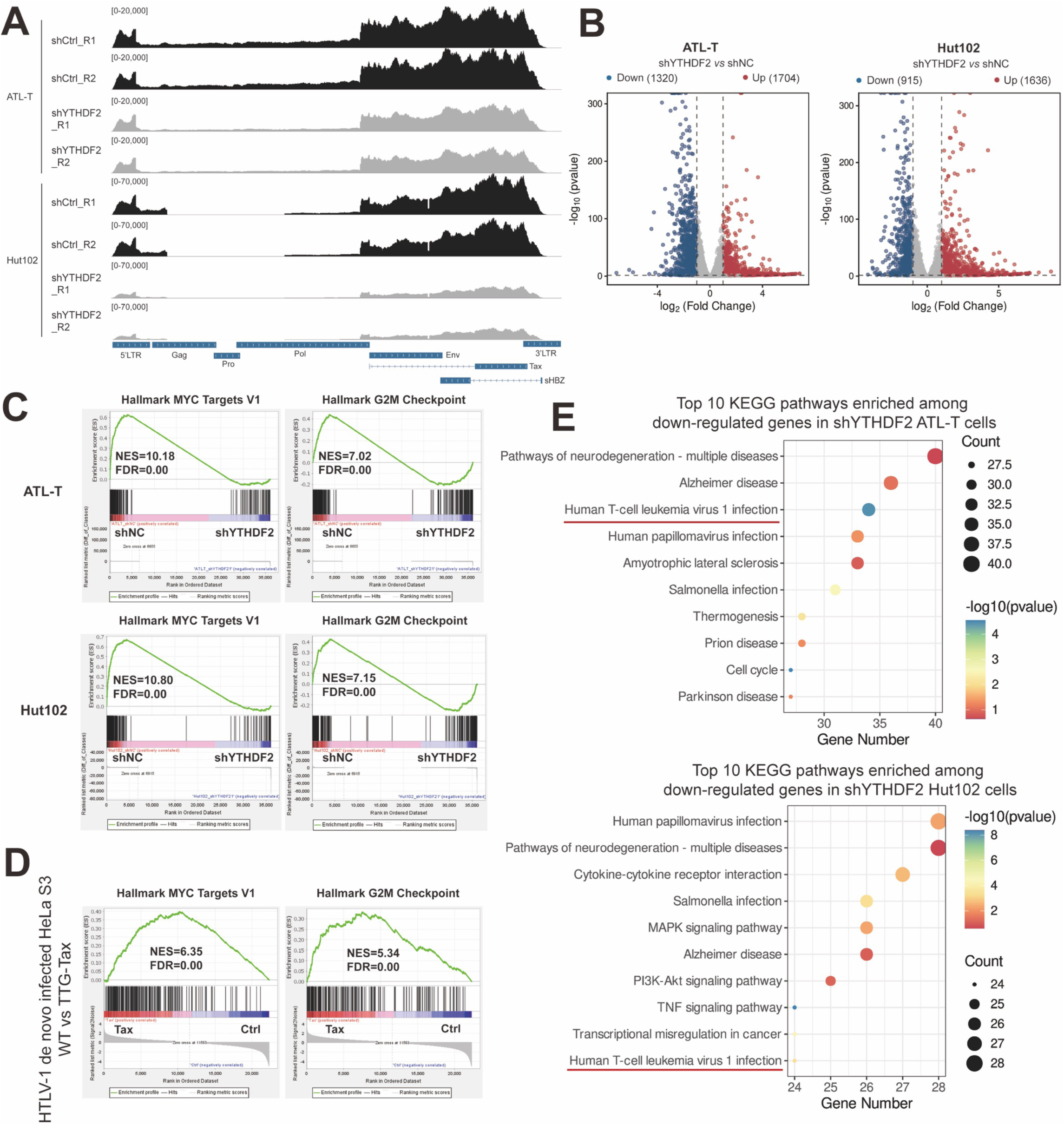
YTHDF2 overlaps with Tax in cellular pathway dysregulation in HTLV-1 infection. (A) RNA-seq reads coverage plot demonstrates a significant reduction in HTLV-1 gene expression levels in (*Upper*) ATL-T and (*Lower*) Hut102 cells following YTHDF2 knockdown (shYTHDF2-1). (B) The volcano plot illustrates differentially expressed genes (Fold change > 1.5, p < 0.05) regulated by YTHDF2 in ATL-T and Hut102 cells. (C) GSEA demonstrating that hallmark genes associated with MYC targets V1 and G2M checkpoint genes sets are downregulated in shYTHDF2 (*Left*) ATL-T and (*Right*) Hut102 cells. FDR < 0.05* for all signatures. (D) GSEA demonstrating that hallmark genes associated with MYC targets V1 and G2M checkpoint genes sets are upregulated in WT vs Tax-KO pX1MT-M transfected HeLa S3 cells. FDR < 0.05* for all signatures. (E) KEGG pathway enrichment of downregulated genes upon YTHDF2 knockdown in (*Upper*) ATL-T and (*Lower*) Hut102 cells.

**Fig. 5.**
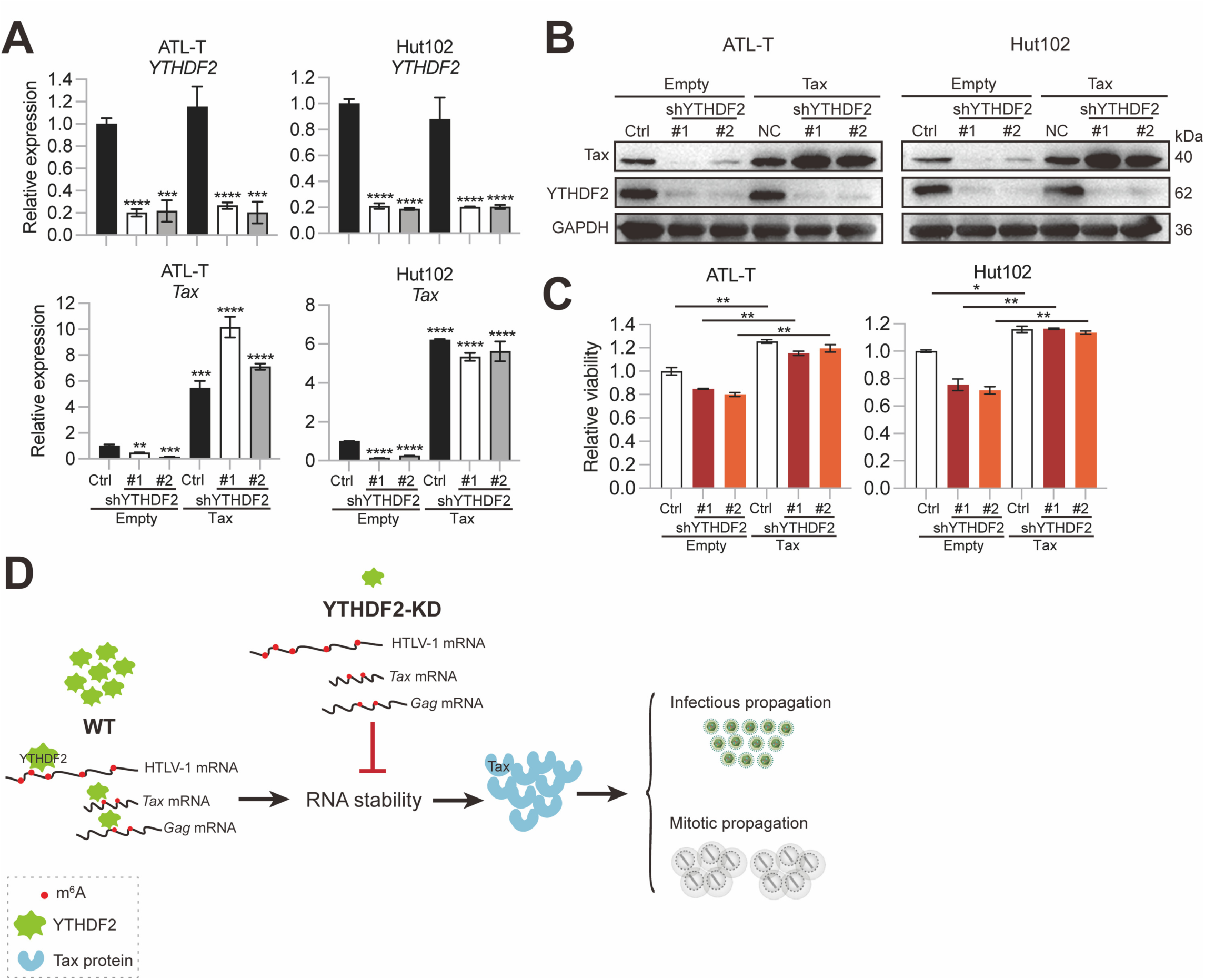
The YTHDF2–m^6^A–Tax axis regulates HTLV-1 persistence. (A) qPCR analysis of YTHDF2 and Tax RNA expression levels in the rescue experiment. (B) Immunoblot analysis of YTHDF2 and Tax protein expression levels in the rescue experiment. (C) CCK-8 assay demonstrates that reintroducing Tax expression in YTHDF2 knockdown cells rescued cell viability. (D) A proposed model illustrating the role of the YTHDF2–m^6^A–Tax axis in HTLV-1 infection. *P* value was calculated using a two-tailed unpaired Student’s t-test. *p<0.05, **p<0.01, ***p<0.001, ****p<0.0001. The results are representatives of three independent experiments.

## Discussion

Our study elucidates for the first time a pivotal role of YTHDF2 in the lifecycle of HTLV-1. The identification of m^6^A modification sites in HTLV-1 RNAs, especially in the pX region that encodes the transactivator gene Tax, underscores the importance of m^6^A modification in HTLV-1 replication. As an m^6^A reader protein, YTHDF2 appears to selectively modulate the stability of various RNAs(28). Although YTHDF2 has been shown to promote RNA decay mostly(28–30), we found that it enhances HTLV-1 RNA stability, consistent with its roles in supporting the infection of other retroviruses such as HIV-1(31, 32). During preparation of this manuscript, we noticed that two other groups (22, 33) have also identified m^6^A modifications in HTLV-1 RNAs . Both King et al. and Gibu et al. used m^6^A inhibitors and demonstrated that m^6^A editing enhances HTLV-1 sense transcription including Tax, which agrees with our observations (22, 33). Different from this study, King et al. analyzed two other m^6^A reader proteins, YTHDF1 and YTHDC1, and showed that YTHDF1 suppresses while YTHDC1 increases HTLV-1 sense transcription(22). These results indicate that the roles of different m^6^A reader proteins in viral infections including HTLV-1 are complex (34); nevertheless, further studies are warranted to address the underlying mechanism in order to better understand the effect of m^6^A modification on HTLV-1 infection.

Mitotic expansion of HTLV-1 infected CD4 T cells contributes enormously to the propagation of HTLV-1 *in vivo*(35). Our findings demonstrate that YTHDF2 supports the proliferation of HTLV-1 infected T cells (Fig. 3), emphasizing its biological significance to the *in vivo* persistence of the virus. The fact that YTHDF2 overlapped with Tax in regulating multiple oncogenic gene sets suggests that YTHDF2 might also contribute to Tax-mediated cellular transformation. On the other hand, YTHDF2 and Tax also have a number of different target genes: further investigation of these genes is expected to better interpret the roles of YTHDF2 in HTLV-1 pathogenesis. In addition, the m^6^A-independent functions of YTHDF2 (36) may also require attention. For example, YTHDF2 is the only YTH reader protein confirmed to recognize 5-methylcytosine (m^5^C), another important posttranscriptional RNA modification involved in the pathogenesis of various diseases(37). It will be interesting to examine in the future whether m^5^C modifications are present in HTLV-1 RNAs and are involved in the YTHDF2–Tax interplay.

The YTH reader proteins are evolutionarily conserved, in particular the three YTHDF proteins; however, each YTHDF protein has a different effect on m^6^A-modified mRNAs(34). For instance, YTHDF1 usually promotes the translation of m^6^A-modified mRNAs through engaging with translation initiation factors, while YTHDF2 generally directs m^6^A-modified mRNAs to decay sites for degradation(34). YTHDF3 can either interact with YTHDF1 to increase the translation of m^6^A-modified mRNAs, or enhance their degradation via binding YTHDF2(34). Unfortunately, it remains still elusive how YTHDF proteins exert different functions despite their high sequence homology. Nevertheless, the opposite effects of YTHDF1 and YTHDC1 on Tax RNA reported by King et al. (22), as well as the Tax RNA-stabilizing function of YTHDF2 shown in this study, highlight the complex roles of m^6^A reader proteins in HTLV-1 infection.

So far we and two other groups (22, 33) independently reported the presence of m^6^A modifications in HTLV-1 RNAs. In addition, we and King et al. showed that at least three m^6^A reader proteins, YTHDF1, YTHDF2, and YTHDC1, can bind m^6^A-edited HTLV-1 RNAs and regulate viral transcription. Yet a key question remains unanswered, which is whether they bind the same or different m^6^A sites in HTLV-1 RNAs. At present it is still debatable as to whether a specific m^6^A residue can recruit specific or all the reader proteins(38). In addition, the location of the m^6^A sites, whether it is in the translated or untranslated regions of viral RNAs, may also lead to distinct outcomes upon recognition by reader proteins(34). Further studies addressing the above points will greatly improve our understanding on the distinct roles of m^6^A reader proteins in HTLV-1 infection.

Overall, this study identifies important m^6^A modification sites in HTLV-1 RNAs and characterizes YTHDF2 as a key m^6^A reader protein required for HTLV-1 infectious and mitotic propagation (Fig. 5D). Given that YTHDF2 has been implicated in tumor development(36), our findings may also shed light on the development of novel targeted therapies for ATLL.

## Materials and Methods

### Cell lines

HTLV-1 infected cell lines, including ATL-T and Hut102, were cultured in RPMI supplemented with 10% fetal bovine serum (FBS). The HTLV-1 uninfected T cell lines, including Jurkat and Hut78, were also cultured in RPMI with 10% FBS. HeLa S3 and HEK293T cells were maintained in DMEM supplemented with 10% FBS. All cell lines were kept in a humidified incubator at 37 °C with 5% CO₂.

### m^6^A modification prediction

For the prediction of m^6^A modification sites in HTLV-1 RNA, the FASTA sequence were input into the SRAMP(23) tool (http://www.cuilab.cn/sramp) using default parameters. High-confidence sites were defined as those with a combined score greater than 0.6.

### MeRIP-qPCR

For m^6^A immunoprecipitation, 200 μg of RNA samples were fragmented using RNA fragmentation buffer (100 mM Tris–HCl, pH 7.0, and 100 mM ZnCl_2_) at 70°C for 10 mins. Standard ethanol precipitation was performed, and 1/10 of the fragmented RNA was retained as the input. The remaining fragmented RNA was immunoprecipitated with anti-m^6^A antibody (56593S, Cell Signaling Technology, USA) in IP buffer (0.1% NP-40, 10 mM Tris-HCl, pH 7.0, 150 mM NaCl) at 4°C for 4 hours. Dynabeads Protein A/G (P2108, Beyotime, China) were washed, added to the mixture, and incubated for 2 hours at 4°C with rotation. The m^6^A-containing RNA fragments were eluted twice with 6.7 mM N6-methyladenosine 5′-monophosphate sodium salt at 4°C for 1 h and precipitated with ethanol at -80°C overnight. The RNA was then dissolved in 10 μL of nuclease-free water and subjected to reverse transcription for qPCR. The primer sequences are listed in Table S1.

### RNA extraction and RT-qPCR

RNA was extracted using the Total RNA Extraction Kit (R701, Vazyme, China), and cDNA was synthesized with the HiScript III 1st Strand cDNA Synthesis Kit (R312, Vazyme, China). The expression levels of the genes of interest were quantified using the QuantStudio 3 Real-Time PCR System (ThermoFisher Scientific, USA). Relative gene expression was quantified using the 2^-ΔΔCt^ method, normalized to *GAPDH*. The primer sequences are listed in Table S2.

### Plasmid construction

Lentiviral vectors (pLKO.1-puro) expressing shRNAs targeting the YTHDF2 coding sequence (CDS) (shYTHDF2-1: CTAGAGAACAACGAGAATAAA or shYTHDF2-2: GCTCCTGGCATGAATACTATA) or a negative control (shNC: CAACAAGATGAAGAGCACCAA) were generated.

HTLV-1 5’ LTR and Tax CDS were amplified from pX1MT-M and cloned into a pME18s backbone (the original promoter SRα is removed), to generate p5’LTR-Tax-WT. Next, the 2nd m^6^A modified base of the Tax CDS in p5LTR-Tax-WT was mutated from A to C to generate p5’LTR-Tax-m^6^A-Mut (Tax amino acid sequence is intact).

The Tax CDS was cloned into the pCSII-CMV-puro-P2A vector to facilitate lentivirus packaging and subsequent infection of T cells, enabling the overexpression of Tax.

### Lentivirus mediated knockdown

A lentivirus-based shRNA system was utilized to knock down YTHDF2 in T cell lines. Plasmids, including pMD2.G (7 μg), psPAX2 (14 μg), and pLKO.1 (14 μg), were transfected into 5.5 × 10^6^ 293T cells in a 10-cm dish using Polyethylenimine (Prime-P100, Serochem, USA) to produce lentivirus. Supernatants were collected at 48 and 72 h, filtered through a 0.45 μm filter, and subsequently used to infect target cells. RNA and protein from the target cells were extracted 48 h post-infection.

### Tax rescue experiment

To overexpress Tax in ATL-T and Hut102 cells, a lentivirus system was utilized. Specifically, 293T cells (5.5×10⁶) were transfected with a mixture of plasmids, including pMD2.G (7 μg), psPAX2 (14 μg), and either pCSII-CMV-puro-P2A or pCSII-CMV-puro-P2A-Tax (14 μg), using polyethylenimine in a 10-cm dish to produce lentivirus. Supernatants were collected at 48 h and 72 h post-transfection and filtered through a 0.45-μm filter. These filtered supernatants were then used to infect cells that had undergone 48 h of knockdown, resulting in Tax overexpression. Cell viability was assessed 48 h post-infection using the CCK-8 assay.

### YTHDF2 siRNA silencing

siRNAs targeting YTHDF2 were synthesized by Gencefe (Wuxi, China). HeLa S3 cells (2 × 10^5) were co-transfected with either specific siYTHDF2 or a non-specific control (siNC, 50 pmol), along with pX1MT-M (1 μg). Supernatants were collected 48 h post-transfection, and total RNA and protein were subsequently extracted from the cells.

### Western blot

Cells were harvested and lysed using RIPA lysis buffer (50 mM Tris-Cl, pH 8; 150 mM NaCl; 2 mM EDTA; 1% Triton X-100; 0.1% SDS), supplemented with a protease inhibitor cocktail (APExBIO, China). Cell lysates were separated by SDS-PAGE, followed by electroblotting onto polyvinylidene difluoride (PVDF) membranes and probing with specific antibodies. The Tax antibody (sc-57872), p19 antibody (sc-57870), and p24 antibody (sc-53891) were obtained from Santa Cruz (USA), while the YTHDF2 (ab220163) and GAPDH (T0004) were purchased from Abcam (USA) and Affinity Biosciences (China), respectively. HRP-linked anti-mouse IgG (7076S) was purchased from Cell Signaling Technology (USA). The WB bands were exposed using ECL luminescent solution (Sparkjade Biotechnology, China) and imaged on the Tanon 5200 Multi (Tanon Life Science, China).

### Enzyme-linked immunosorbent assay (ELISA)

Supernatants from cultured cells were centrifuged at 1,710 × g for 5 min to remove debris and then diluted and quantified for p19 by ELISA (ZeptoMetrix, USA) according to manufacturer’s instructions.

### RIP-qPCR

2×10^7^ cells were collected using a cell lysate containing protease and RNase inhibitors (20 mM Tris-HCl pH 7.4, 100 mM KCl, 5 mM MgCl_2_, 0.5% Triton X-100). Magnetic beads (BeyoMag Protein A+G) were pre-bound with either Rabbit IgG or anti-YTHDF2, followed by incubation with the cell lysate in immunoprecipitation buffer (50 mM Tris-HCl pH 7.4, 150 mM NaCl, 1 mM MgCl2, 0.05% Triton X-100, 20 mM EDTA pH 8.0, 1 mM DTT, 0.2 U/µL RNase inhibitor) overnight at 4°C. The magnetic bead complexes were washed and digested with proteinase K digestion buffer for 30 minutes at 55°C, and RNA was extracted using TRIzol. Specific fragment enrichment was assessed by RT-qPCR, with relative enrichment normalized to input samples, calculated as follows: ΔCt[normalized RIP]=(Average Ct[RIP]−(Average Ct[Input]−log2(Input Dilution Factor))), %Input=2^−ΔCt[normalizedRIP]^(39).

### RNA stability assay

ATL-T and Hut102 cells were treated with 5 µg/mL actinomycin D (HY-17559, MedChemExpress, USA) for 0, 4, 8, and 12 h, followed by total RNA extraction using TRIzol. RT-qPCR was performed to analyze mRNA levels, with specific primers listed in Table S2. The mRNA half-life (t_1/2_) was calculated using the equation: t_1/2_ = ln 2/k_decay_.

### Cell Counting Kit-8 (CCK-8) analysis

100 μL of the cell suspension was transferred to a 96-well plate, and 10 μL of CCK-8 reagent (Adamas Life, China) was added. The cells were then incubated in the dark in a humidified incubator at 37°C with 5% CO₂. After 3 h, the absorbance at 450 nm was measured using a plate reader (Berthold, Germany).

### Cell apoptosis analyses

Cell apoptosis was detected using a cell apoptosis detection kit (E-CK-A211, Elabscience, China) according to the manufacturer’s instructions. Specifically, cells were collected and stained with FITC Annexin V and PI at room temperature for 15 minutes. Following this, a binding buffer was added to mix the samples, which were then analyzed using a NovoCyte flow cytometer (Agilent Technologies, USA).

### RNA-seq and bioinformatic analyses

HeLa S3 cells were seeded at a density of 2×10⁵ cells/ml in a 12-well plate, and the cells were transfected with 1 μg of pX1MT-M-WT or TTG-Tax(24) using Lipofectamine 2000 (Thermo). After 48 hours, total RNA was extracted.

For RNA-seq analysis of HTLV-1 *de novo* infected HeLa S3 cells and HTLV-1 infected cells with YTHDF2 knockdown, library preparations were conducted using a mix of 250 ng total RNA and 0.25 ng Spike-in RNA Variant Control Mix E2 (Lexogen, Austria) with the TruSeq RNA Library Prep Kit (Illumina). The libraries were quantified by qPCR using the KAPA Library Quantification Kit for Illumina Libraries (Kapa Biosystems, USA), and library profiles were evaluated using an Agilent 2100 Bioanalyzer (Agilent Technologies, USA).

For raw reads of RNA-seq, alignment was conducted with STAR v2.7.11a(40). The unique alignments were extracted with Samtools v1.19(41). To identify DEGs, reads were counted using FeatureCount v2.22.0(42), and DESeq2 v1.46.0(43) was applied with the following thresholds: FC ≥ 2 and p-value < 0.05. Gene Set Enrichment Analysis (GSEA) v4.3.2 was conducted for pathway regulation analysis.

### Animal experiments

4 to 5 weeks old female NSG mice (Hangzhou Ziyuan Experimental Animal Technology Co., Ltd., China) were maintained in a specific pathogen-free room at the Animal Experimental Center of China Pharmaceutical University. 5.0×10^6^ control or YTHDF2 knockdown Hut102 cells were resuspended in 100 µL of precooling PBS and subcutaneously injected into mice. At the end of the experiment, mice were sacrificed, and tumors were collected for further measurements or analyses. Tumor size is estimated by measuring the longest diameter of the entire tumor and the corresponding vertical diameter (Tumor volume (mm^3^) = 0.5 × L(length) × W^2^ (width)). All animal experiments were approved by the Institutional Animal Care Board of China Pharmaceutical University.

### Data availability

The RNA-Seq data reported in this paper have been deposited in the Sequence Read Archive database, https://www.ncbi.nlm.nih.gov/sra (accession no. PRJNA1331476).

### Statistical analyses

Data analysis and plotting were done using R 4.4.2 and GraphPad Prism 8. Statistical analyses were performed by using the unpaired Student t-test. A value of *P* < 0.05 was considered statistically significant.

## Supplement Figures

**Fig. S1.**
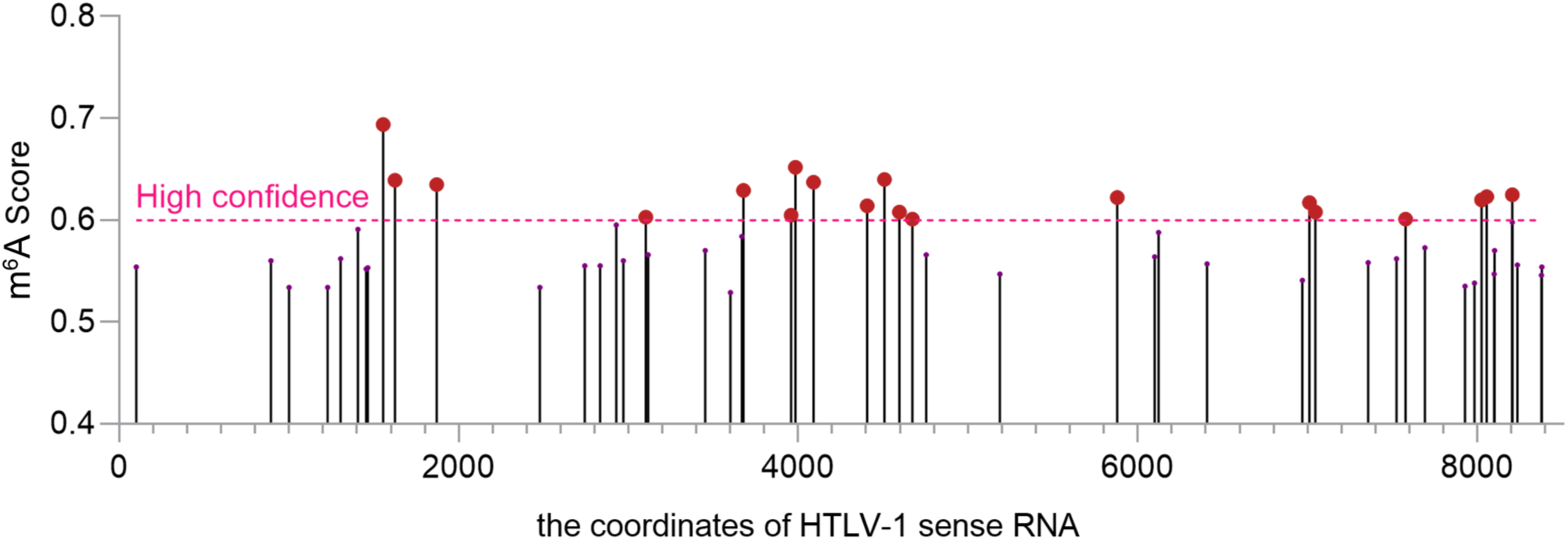
Prediction of m^6^A modification sites on HTLV-1 RNA. Using the SRAMP tool, 19 high-confidence m^6^A modification sites were predicted on the HTLV-1 RNA.

**Fig. S2.**
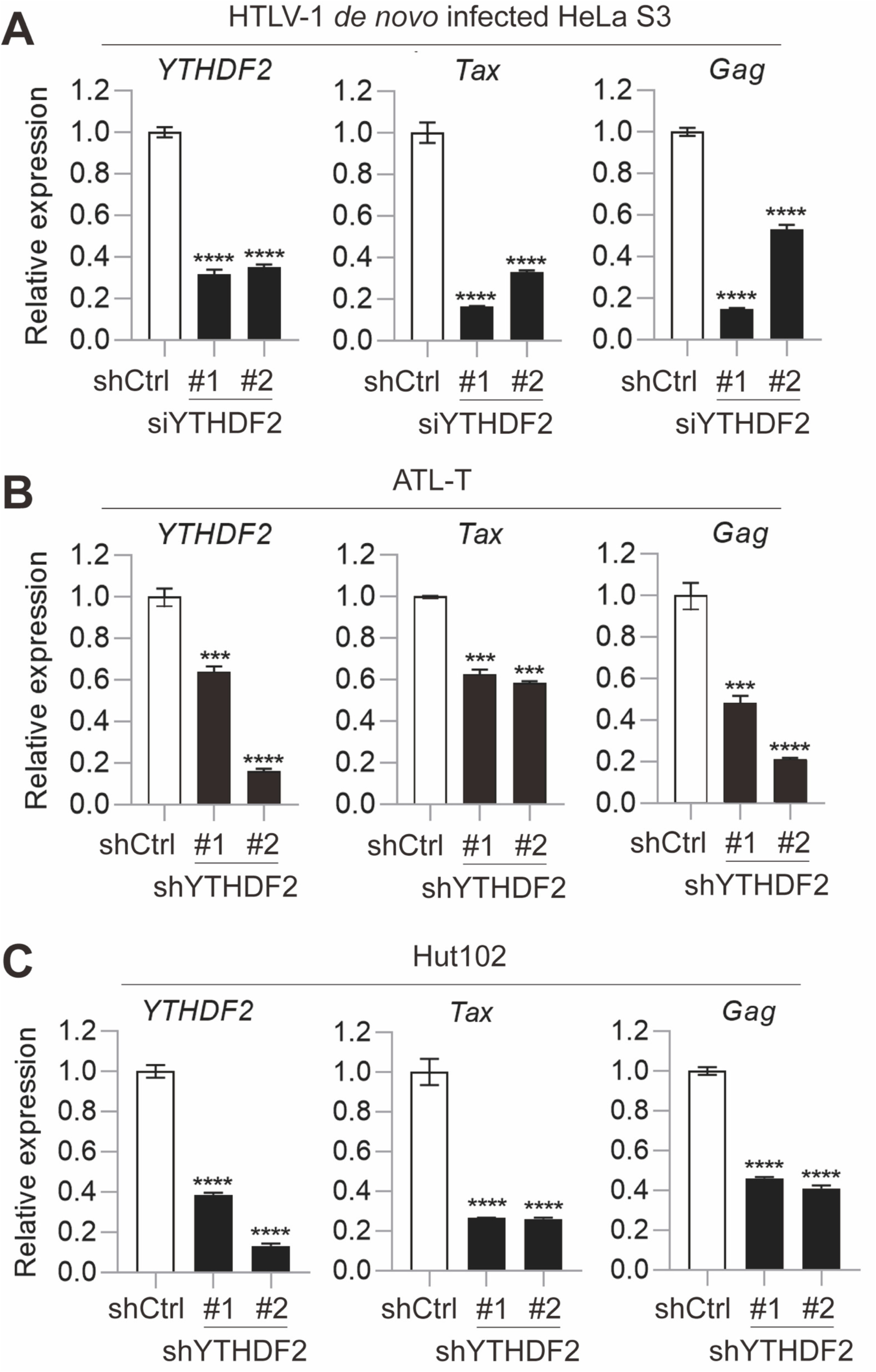
YTHDF2 knockdown suppresses HTLV-1 Tax and gag RNA expression. Following YTHDF2 knockdown via siRNA in (A) HeLa S3 or shRNA in (B) ATLT and (C) Hut102 cells, qPCR was used to quantify the expression levels of YTHDF2, Tax, and Gag.

**Fig. S3.**
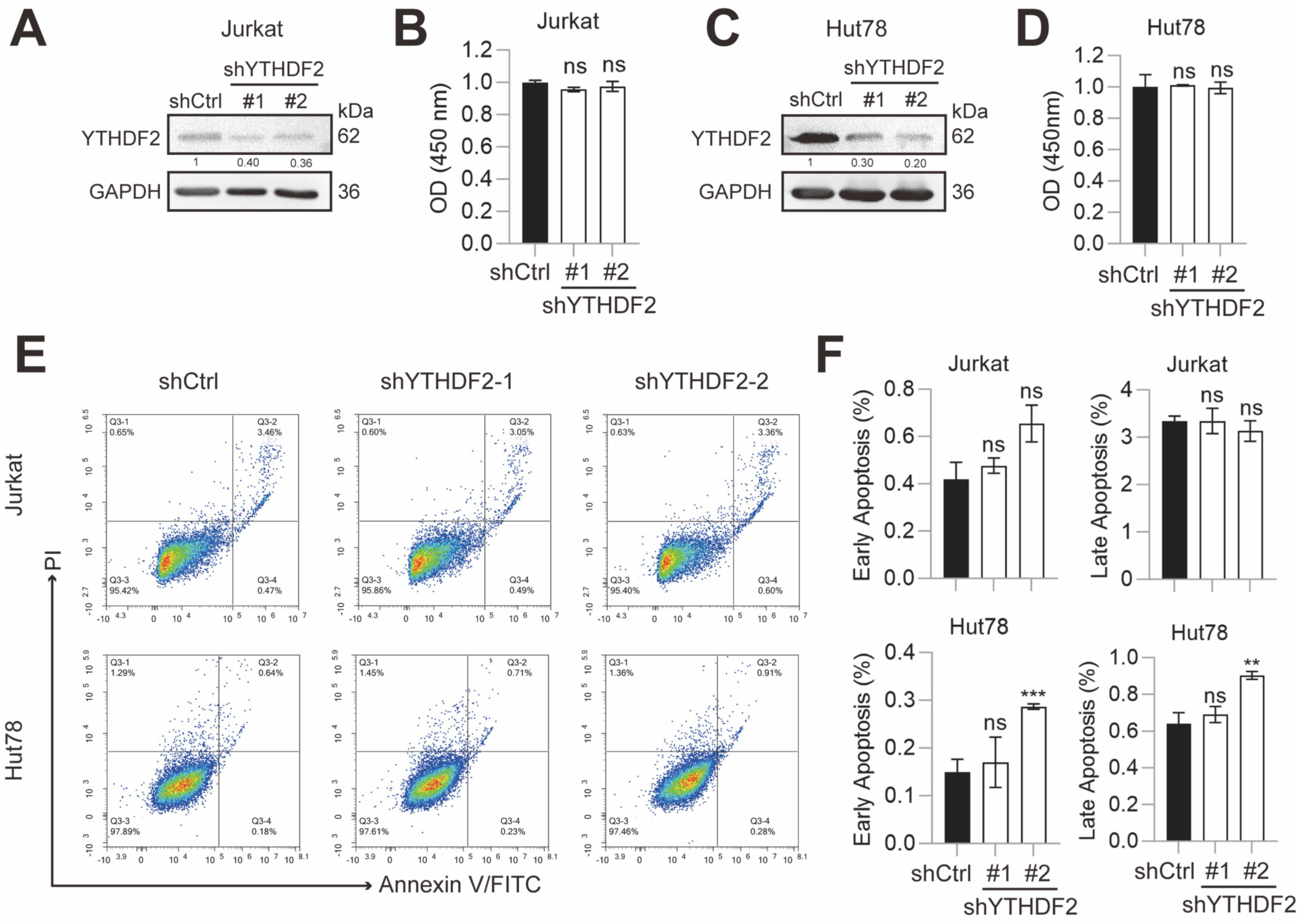
Effects of YTHDF2 knockdown on proliferation and apoptosis in HTLV-1 negative Jurkat and Hut78 cells. (A and C) Immunoblot analysis of YTHDF2 knockdown efficiency in (A) Jurkat and (C) Hut78 cells, with GAPDH used for normalization. (B and D) Cell proliferation was measured by CCK-8 assay at 72 h after YTHDF2 knockdown in (B) Jurkat and (D) Hut78 cells. (E) Flow cytometry analysis of apoptosis in (*Upper*) Jurkat and (*Lower*) Hut78 cells after YTHDF2 knockdown. (F) Statistical analysis of early and late apoptosis based on the data in (E). *P* value was calculated using a two-tailed unpaired Student’s t-test. **p<0.01, ***p<0.00. ns, not significant. The results are representatives of three independent experiments.

**Fig. S4.**
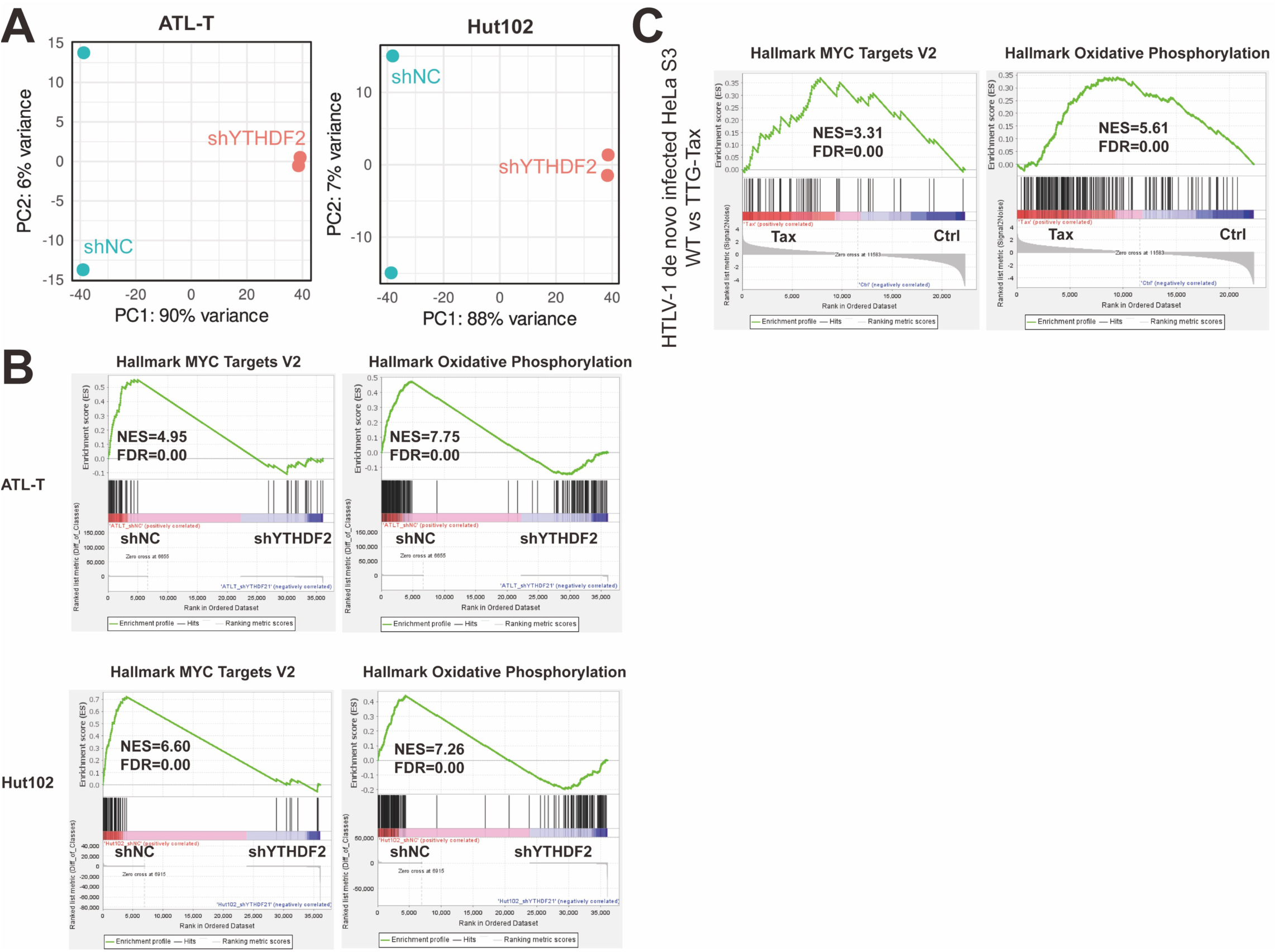
Transcriptomic results show YTHDF2 and Tax regulate overlapping pathways. (A) PCA of RNA-seq from YTHDF2 knockdown ATL-T and Hut102 cells. (B) GSEA demonstrating that hallmark genes associated with MYC targets V2 and Oxidative phosphorylation genes sets are downregulated in shYTHDF2 ATL-T and Hut102 cells. FDR < 0.05* for all signatures. (C) GSEA demonstrating that hallmark genes associated with MYC targets V2 and Oxidative phosphorylation genes sets are upregulated in WT vs Tax-KO pX1MT-M transfected HeLa S3 cells. FDR < 0.05* for all signatures.

## Acknowledgements

This study was supported by the National Natural Science Foundation of China (32270171 and 32070155 to G. M.). The funders had no role in study design, data collection and interpretation, or the decision to submit the work for publication.

